# A single PLAT domain protein couples reproductive arrest and carotenoid pigmentation during diapause in the two-spotted spider mite, *Tetranychus urticae* Koch

**DOI:** 10.64898/2026.05.13.724795

**Authors:** Rismayani, Kanae Sai, Tomohiro Ohsako, Murata Kohyoh, Yuka Arai, Naoki Takeda, Masanobu Yamamoto, Rika Umemiya-Shirafuji, Takeshi Suzuki

## Abstract

Adult females of the two-spotted spider mite, *Tetranychus urticae* Koch, enter a photoperiodically induced diapause to overwinter. Diapause in *T. urticae* is accompanied by reproductive arrest and the orange body coloration that arises from the accumulation of astaxanthin esters. How these two traits are coordinated at the molecular level remains poorly understood. Here, we compared the proteomes of adult females reared under diapause-inducing (long-night) and non-diapause-inducing (short-night) photoperiods using liquid chromatography–tandem mass spectrometry, followed by RNA interference (RNAi) of candidate genes. The carotenoid biosynthesis enzymes phytoene desaturase (TuPDS) and lycopene cyclase/phytoene synthase (TuLCPS), both encoded by genes horizontally transferred from fungi, were more abundant in diapausing females than in non-diapausing females. RNAi of the genes encoding TuPDS and TuLCPS markedly reduced orange pigmentation as well as β-carotene and astaxanthin contents, demonstrating that these enzymes are required for diapause-associated pigmentation. Our proteomic analysis further identified a single PLAT (Polycystin-1, Lipoxygenase, Alpha-toxin) domain protein, TuPLAT10, as one of the most strongly upregulated proteins in diapausing females. The PLAT domain is a lipid-binding module, suggesting a role for TuPLAT10 in lipid metabolism. In addition to the suppression of orange pigmentation, RNAi of the *TuPLAT10* gene restored reproduction even under diapause-inducing conditions and selectively reduced TuPDS and TuLCPS protein levels, despite the absence of sequence similarity to their genes. We propose that TuPLAT10 acts as a lipid-allocation switch that, in response to photoperiodic information, partitions fatty acids between astaxanthin esterification and yolk lipid supply, thereby coupling reproductive arrest and carotenoid pigmentation during diapause in *T. urticae*.

## 1. Introduction

Diapause is a state of developmental arrest that enables arthropods to survive unfavorable seasons and synchronize their life cycles (Tauber et al., 1986). Obligate diapause occurs constitutively in every generation, whereas facultative diapause is induced by environmental cues, of which photoperiod is the most reliable predictor of seasonal change and is integrated during a photoperiodic sensitive stage (Goto, 2016).

The two-spotted spider mite, *Tetranychus urticae* Koch (Trombidiformes: Tetranychidae), is a globally distributed arthropod herbivore and one of the most economically important agricultural and horticultural pests, feeding on more than 1,100 plant species across more than 140 plant families (Grbić et al., 2011; Migeon and Dorkeld, 2025). In temperate environments, adult *T. urticae* females enter a facultative reproductive diapause following exposure to long-night photoperiods during the immature stages (Veerman, 1985; Suzuki and Takeda, 2009). Diapause in *T. urticae* females is marked by a suite of physiological and morphological adaptations, most notably the arrest of reproduction and a striking change in body pigmentation from yellow-green to orange (Veerman, 1974). This body color change is attributed to the accumulation of astaxanthin, a ketocarotenoid that exists predominantly as fatty acid esters in diapausing *T. urticae* females (Kawaguchi et al., 2016) and protects against multiple environmental stresses (Suzuki et al., 2009, 2015; Ghazy and Suzuki, 2014).

Carotenoids are terpenoid-based pigments widely distributed in nature. In animals, they fulfill essential roles in pigmentation, photoreception, oxidative stress protection, and signaling (von Lintig et al., 2005; Heath et al., 2013). Although animals were long considered incapable of synthesizing carotenoids *de novo*, this view changed when horizontal gene transfer (HGT) of fungal carotenoid biosynthesis genes was discovered in arthropods, including aphids, gall midges, and spider mites (Moran and Jarvik, 2010; Grbić et al., 2011; Cobbs et al., 2013). In *T. urticae*, the HGT-derived carotenoid biosynthesis genes *phytoene desaturas*e (*TuPDS*) and *lycopene cyclase/phytoene synthase* (*TuLCPS*) underlie the endogenous synthesis of β-carotene (Altincicek et al., 2012). In addition, a cytochrome P450 of the CYP3 clan, CYP384A1, was implicated by genetic mapping as a candidate carotenoid ketolase associated with astaxanthin synthesis from β-carotene in the closely related species *Tetranychus kanzawai* Kishida (Wybouw et al., 2019). Genetic analyses of albino mutants implicated β-carotene or its derivatives (e.g., retinal) as photoreceptor pigments required for photoperiodic time measurement that drives diapause induction in *T. urticae* (Veerman and Helle, 1978; Veerman, 1980). Using bulked segregant analysis, Bryon et al. (2017) identified the albino locus as the *TuPDS* gene, thus providing a molecular basis for these earlier genetic findings. In addition to such roles in pigmentation and photoreception, the predominance of astaxanthin as fatty acid esters in diapausing females (Kawaguchi et al., 2016), together with the role of triacylglycerols and phospholipids as essential lipid components for oogenesis, points to a central role of lipid metabolism in coordinating carotenoid-based pigmentation and reproductive arrest in *T. urticae*.

Previous transcriptomic and proteomic studies revealed that a broad range of genes are differentially expressed during diapause in *T. urticae*, including upregulation of carotenoid biosynthesis genes as well as genes involved in Ca^2+^ signaling, cytoskeletal reorganization, and metabolic suppression (Bryon et al., 2013; Zhao et al., 2017). However, the direct functional contributions of individual carotenoid biosynthesis genes to diapause-associated pigmentation have not been demonstrated. In addition, the regulators linking lipid metabolism to carotenoid biosynthesis and reproductive arrest have yet to be identified. To address these gaps, we used liquid chromatography–tandem mass spectrometry (LC-MS/MS) to perform comparative proteome analysis between adult *T. urticae* females reared under diapause-inducing and non-diapause-inducing conditions from the immature stages, followed by RNAi-mediated functional analysis of candidate genes. This approach allowed us to identify the carotenoid biosynthesis enzymes required for diapause-associated pigmentation and revealed a single PLAT (Polycystin-1, Lipoxygenase, Alpha-toxin) domain protein, TuPLAT10, as a novel regulator that couples reproductive arrest and carotenoid pigmentation in *T. urticae*.

## 2. Materials and methods

### 2.1. Mite rearing

The *T. urticae* population was collected from apple trees (*Malus pumila* Mill. ‘Fuji’) in Akita, Japan. The mites were reared in the laboratory on primary leaves of kidney bean (*Phaseolus vulgaris* L.) under a short-night photoperiod of L:D 16:8 at an air temperature of 25 °C. Adult *T. urticae* females of random ages were collected using an air pump–based system (Cazaux et al., 2014), transferred onto kidney bean leaves placed in Petri dishes, and allowed to lay eggs for 24 h. All adult females were then removed, leaving only the eggs. The dishes were covered with lids and sealed with stretched Parafilm M (Bemis NA, Neenah, WI, USA) to maintain approximately 100% relative humidity (RH) inside the dishes to synchronize hatching (Suzuki et al., 2017a). The sealed dishes were placed in an incubator (MIR-154; PHC Corp., Tokyo, Japan) set at 25 °C under a L:D 16:8 photoperiod. After 3 days, the lids were opened and the incubated eggs were exposed to ∼50% RH for at least 2 h to stimulate hatching. Newly hatched larvae were reared in an incubator set at 18 °C under L:D 16:8 or L:D 8:16 photoperiod to induce reproduction (non-diapause) or diapause at the adult stage, respectively.

### 2.2. LC-MS/MS

A total of 100 females reared under L:D 16:8 or L:D 8:16 were collected daily from day 0 to day 7 after adult emergence and transferred into a 1.5-mL polypropylene tube (100 mites/tube). Samples were collected in three biological replicates per day for each condition, yielding a total of 48 tubes (2 conditions × 8 days × 3 replicates). The protocol for protein extraction, digestion, and peptide desalting followed Arai et al. (2025) and Takeda et al. (2026), with brief descriptions provided below.

For protein extraction, protein lysis buffer containing 10 mM Tris-HCl (pH 9.0) and 8 M urea was added to the tubes. The mites were homogenized using a pestle and an ultrasonicator (AS38A; As One, Osaka, Japan). Homogenized samples were centrifuged at 13,760 ×*g* (5424 R; Eppendorf AG, Hamburg, Germany) for 10 min at 4 °C to remove cell debris, and the supernatant was collected for further steps. Protein concentration was measured using a Qubit 4 fluorometer (Thermo Fisher Scientific, Waltham, MA, USA). The extracted proteins were reduced with 10 mM dithiothreitol (Invitrogen, Waltham, MA, USA) for 30 min and then alkylated with 50 mM iodoacetamide (Fujifilm Wako Pure Chemical, Osaka, Japan) for 20 min in the dark. The solution was diluted with 50 mM NH_4_HCO_3_ (Fujifilm Wako Pure Chemical) to a quarter of the original urea concentration, and proteins were digested with trypsin (trypsin:protein = 1:50; Promega K.K., Tokyo, Japan) at 25 °C for 12 h. The reaction was stopped by addition of 2% trifluoroacetic acid (TFA) in the same volume as the sample solution, followed by centrifugation at 13,760 ×*g* at 4 °C for 10 min. The peptide-desalting column (200 µL; GL-Tip SDB; GL Sciences Inc., Tokyo, Japan) was activated with solution A (water:acetonitrile:TFA = 200:800:1) and equilibrated with solution B (water:acetonitrile:TFA = 950:50:1) by centrifugation at 1,100 ×*g* at 4 °C for 3 min each. The samples were then loaded onto the column, washed with solution B, and the digested peptides were eluted with solution A. The eluted samples were concentrated in a centrifugal vacuum evaporator consisting of a centrifugal concentrator (CC-105; Tomy Seiko Co., Ltd., Tokyo, Japan), a cold trap (TU-500; Tomy Seiko Co., Ltd.), and a diaphragm vacuum pump (DTU-20; ULVAC, Kanagawa, Japan). Concentrated samples were then stored at −80 °C until LC-MS/MS analysis.

Shotgun proteomic analysis was conducted using an Orbitrap Exploris 480 (Thermo Fisher Scientific). The raw mass spectrometry data were deposited at jPOSTrepo (https://repository.jpostdb.org/; accession JPST004604; Okuda et al., 2017) and ProteomeXchange (https://www.proteomexchange.org/; accession PXD078207; Deutsch et al., 2023). Mass spectrometry data were analyzed using Proteome Discoverer v.2.5.0.400 software (Thermo Fisher Scientific). The protein database was annotated using the *T. urticae* proteome (v.20190125) from the ORCAE database (http://bioinformatics.psb.ugent.be/orcae/overview/Tetur; Sterck et al., 2012). The search parameters were as follows: trypsin was specified as the enzyme with a maximum of two missed cleavage sites; precursor mass tolerance was set to 20 ppm and fragment mass tolerance to 0.05 Da; and the false discovery rate was set to 1%. Differentially expressed proteins were identified based on a fold change of ≥2 and a *p*-value of <0.05.

### 2.3. RNAi

The nucleotide sequences of *TuPDS* (*tetur01g11270*), *TuLCPS* (*tetur01g11260*), *TuCYP384A1* (*tetur38g00650*), and *TuPLAT10* (*tetur11g05720*) were obtained from the ORCAE database. Total RNA was extracted from frozen adult females using NucleoSpin RNA Plus XS (Macherey-Nagel GmbH & Co. KG, Düren, Germany) according to the manufacturer’s protocol. The concentration of the extracted RNA was measured with a spectrophotometer (NanoPhotometer N60; Implen, Munich, Germany). cDNA was reverse-transcribed from the extracted total RNA using a SuperScript II cDNA Synthesis Kit (Thermo Fisher Scientific) and stored at −30 °C. Genomic DNA (gDNA) was extracted using a NucleoSpin Tissue Extraction Kit (Macherey-Nagel) and stored at −30 °C. Using cDNA as a template, a 625-bp fragment of *TuPDS*, a 635-bp fragment of *TuLCPS*, a 649-bp fragment of *TuCYP384A1*, and a 325-bp fragment of *TuPLAT10* were PCR-amplified using KOD-Plus DNA polymerase (Toyobo, Osaka, Japan). A 382-bp intergenic fragment (non-coding [NC], genomic coordinate: scaffold 12, position 1690614–1690995; Suzuki et al., 2017b) was PCR-amplified in the same manner using gDNA as a template. Primers used to amplify the DNA fragments of *TuPDS*, *TuLCPS*, *TuCYP384A1*, *TuPLAT10*, and NC are listed in Table S1. Amplified DNA fragments were purified using a NucleoSpin Gel and PCR Clean-Up Kit (Macherey-Nagel). RNA fragments were transcribed from each DNA template using an *in vitro* Transcription T7 Kit (Takara Bio, Kusatsu, Japan). After treatment with DNase I (Takara Bio) for 30 min, the RNA fragments were denatured at 95 °C for 5 min and slowly cooled to 25 °C to facilitate the formation of dsRNA. The dsRNAs were then purified by phenol-chloroform extraction and ethanol precipitation.

A paraffin wax film (Parafilm M; Bemis, Neenah, WI, USA) folded in four was placed at the bottom of a plastic Petri dish, and the four corners were crimped. A piece of Kimwipe placed on the film was soaked with 10 µL of dsRNA solution at 1 µg µL⁻¹ containing 0.1% (v/v) Tween 20 and 0.1% (w/v) Brilliant Blue FCF (Fujifilm Wako Pure Chemical). Following Bensoussan et al. (2020), mites were placed ventral side down on the dsRNA-soaked Kimwipe and incubated at 25 °C for 2 h. In the present study, RNAi treatment was applied twice to each individual: first to newly hatched larvae and then to newly emerged adults. Because the cuticle of *T. urticae* is semi-transparent, color changes in the midgut can be assessed externally under a stereomicroscope (Bensoussan et al., 2018). Individuals whose midgut had turned blue, indicating successful ingestion of the dsRNA solution, were selected for evaluation of RNAi effects in the following experiments.

### 2.4. Body color evaluation

After oral delivery of dsRNAs targeting *TuPDS*, *TuLCPS*, *TuCYP384A1*, *TuPLAT10*, and NC (ds*TuPDS*, ds*TuLCPS*, ds*TuCYP384A1*, ds*TuPLAT10*, and dsNC, respectively), the mites were transferred onto bean leaves and reared in an incubator set at 18 °C under L:D 8:16 for 7 days. Adult females were then mounted in a solution of 50% (v/v) glycerol diluted in 1× PBS and 0.1% (v/v) Tween 20 on a glass slide, and their dorsal sides were photographed under bright field with a color CMOS camera (K7; Leica Microsystems, Wetzlar, Germany) connected to a stereomicroscope (M205 FA; Leica Microsystems). Using ImageJ (Schneider et al., 2012), we extracted a 200 × 850-pixel image region from the dorsal surface of each female, immediately posterior to the eye spots, and obtained its RGB values. The RGB values of each individual were converted to normalized chromaticity coordinates using the following formulas (Stevens et al., 2007; Troscianko and Stevens, 2015): RG = (R − G) / (R + G) and GB = (G − B) / (G + B). Based on these values, the body color of each female was classified into one of three categories: orange (0.15 < RG < 0.60 and −0.10 < GB < 0.20), yellow (−0.05 < RG < 0.15 and 0.10 < GB < 0.50), or green (−0.36 < RG < −0.05 and 0.31 < GB < 0.98).

### 2.5. High-performance liquid chromatography

Following the same dsRNA treatment and rearing procedure described in section 2.4, adult females from each RNAi treatment were classified by body coloration phenotype (orange, yellow, or green; see section 2.4), and the subgroups containing sufficient individuals (*n* = 100) were collected using an air pump–based system (Cazaux et al., 2014). The collected mites were homogenized using a polypropylene pestle homogenizer in a 1.5-mL polypropylene tube. Then 300 µL of acetone was added to the mite sample, and the mixture was sonicated for 3 min to aid in extraction. The homogenates were transferred to a spin column tube equipped with a 0.22-µm-pore filter (Ultrafree-MC; Merck, Darmstadt, Germany), and debris was removed by centrifugation at 12,000 ×*g* for 4 min.

Each extract was aliquoted into two tubes (50 µL/tube), each prefilled with 200 µL of 50 mM Tris-HCl buffer (pH 7.5). Because astaxanthin accumulates predominantly as fatty acid esters during diapause in *T. urticae* (Kawaguchi et al., 2016), cholesterol esterase (CE) treatment was performed to hydrolyze the esters and enable quantification of total astaxanthin as the free form. To one tube (CE-digested sample), 20 µL of the Tris-HCl buffer containing CE (1 unit/µL) was added, whereas 20 µL of the Tris-HCl buffer without CE was added to the other tube as a control (undigested sample). Both samples were incubated at 37 °C for 2 h. After incubation, 100 mg of sodium sulfate hydrate was added to each sample, followed by 200 µL of petroleum ether. The samples were vortexed for 30 s and centrifuged at 1,100 ×*g* for 3 min to separate the ether and aqueous layers. The aqueous phase was used for total protein quantification with the Qubit 4 fluorometer. The ether phase, which contained the carotenoids, was collected and transferred to 1.5-mL tubes containing anhydrous sodium sulfate crystals to dehydrate. The remaining aqueous phase was re-extracted three times with petroleum ether, and the ether phases were pooled. The combined ether phase with the sodium sulfate crystals was centrifuged at 1,250 ×*g* for 3 min, and the supernatant was collected. The remaining sodium sulfate crystals were washed three times with petroleum ether, and the washings were combined with the supernatant. The combined ether fraction was dried under a nitrogen stream, dissolved in 50 µL of acetone and stored at −25 °C in the dark until high-performance liquid chromatography (HPLC) analysis. The undigested samples (−CE) were used to quantify β-carotene and free astaxanthin, whereas the CE-digested samples (+CE) were used to quantify total (free plus esterified) astaxanthin.

For HPLC analysis, standard β-carotene (Nacalai Tesque, Inc., Kyoto, Japan) and astaxanthin (Dr. Ehrenstorfer GmbH, Augsburg, Germany) were dissolved in chloroform to prepare stock solutions of each standard at 10 mg mL^−1^. The stock solutions were diluted 1,000-fold, 10,000-fold, and 100,000-fold with methanol to yield working standard solutions at final concentrations of 10, 1, and 0.1 µg mL^−1^, respectively. Using a glass microsyringe (GL Sciences, Tokyo, Japan), 5 µL of each sample extract and each working standard solution was injected into an HPLC system (PU-980; Jasco, Tokyo, Japan) equipped with a UV/visible detector (UV-970 Intelligent UV/VIS Detector; Jasco) and ODS column (Wakosil-II 5C18 AR 4.6 mm × 250 mm; Fujifilm Wako Pure Chemical). β-carotene and astaxanthin were quantified by gradient elution using acetone and MilliQ water as the mobile phases, with the flow rate set at 0.8 mL min^−1^ and the detection wavelength at 478 nm.

### 2.6. Mite performance assay

After oral delivery of ds*TuPDS*, ds*TuLCPS*, ds*TuCYP384A1*, ds*TuPLAT10*, and dsNC, mites were transferred onto bean leaf discs (10 mm diameter; 1 mite/disc) placed on water-soaked cotton in a Petri dish. These mites were placed in the incubator set at 18 °C under L:D 8:16. Mite fecundity and survival were observed daily for 10 and 12 days, respectively. Leaf discs were replaced with fresh ones every 3 days.

### 2.7. Statistical analysis

All data were analyzed using R v.4.2.2. Differences in relative protein abundances of TuPDS, TuLCPS, TuCYP384A1, and TuPLAT10 between diapausing and non-diapausing adult females were analyzed by using Student’s *t*-test (*t.test* function in the R package *stats*). The proportions of body color phenotypes (orange, yellow, and green) among dsRNA treatments were compared with the negative control (dsNC) by applying Fisher’s exact test (*fisher.test* function in the R package *stats*). Total β-carotene and astaxanthin contents in dsRNA-treated mites (CE-digested and undigested samples) were compared with dsNC by using Dunnett’s test (*glht* function in the R package *multcomp*). Egg production per female per day and protein abundances among dsRNA treatments were analyzed by performing Dunn’s multiple comparison test (*dunnTest* function in the R package *FSA*). The survival of adult females after RNAi-mediated gene silencing was plotted using the Kaplan–Meier method (*survfit* function in the R package *survival*), and differences in survival curves among treatments were analyzed by applying the log-rank test with Bonferroni correction (*survdiff* function in the R package *survival*).

## 3. Results

### 3.1. Dynamic proteomic changes during the diapause-associated body color transition in *T. urticae*

Diapausing and non-diapausing *T. urticae* females showed divergent body pigmentation patterns over the 7-day observation period following the final molt to adulthood (Fig. 1A). On day 0, females from both groups were pale and visually indistinguishable. From day 1 onward, diapausing females developed a uniform orange pigmentation that progressively intensified throughout the time course. In parallel, the dark-colored digestive cell aggregates initially observed in the ventriculus and caeca shifted toward the posterior midgut by day 4 and were subsequently excreted, such that by day 7 these aggregates were no longer visible and the entire body displayed an orange pigmentation. In contrast, non-diapausing females gradually acquired a yellowish green pigmentation over the same period, with the dark-colored digestive cell aggregates in the ventriculus and caeca becoming progressively more distinct. Body size also increased in non-diapausing females, accompanying ovarian development.

**Fig. 1.**
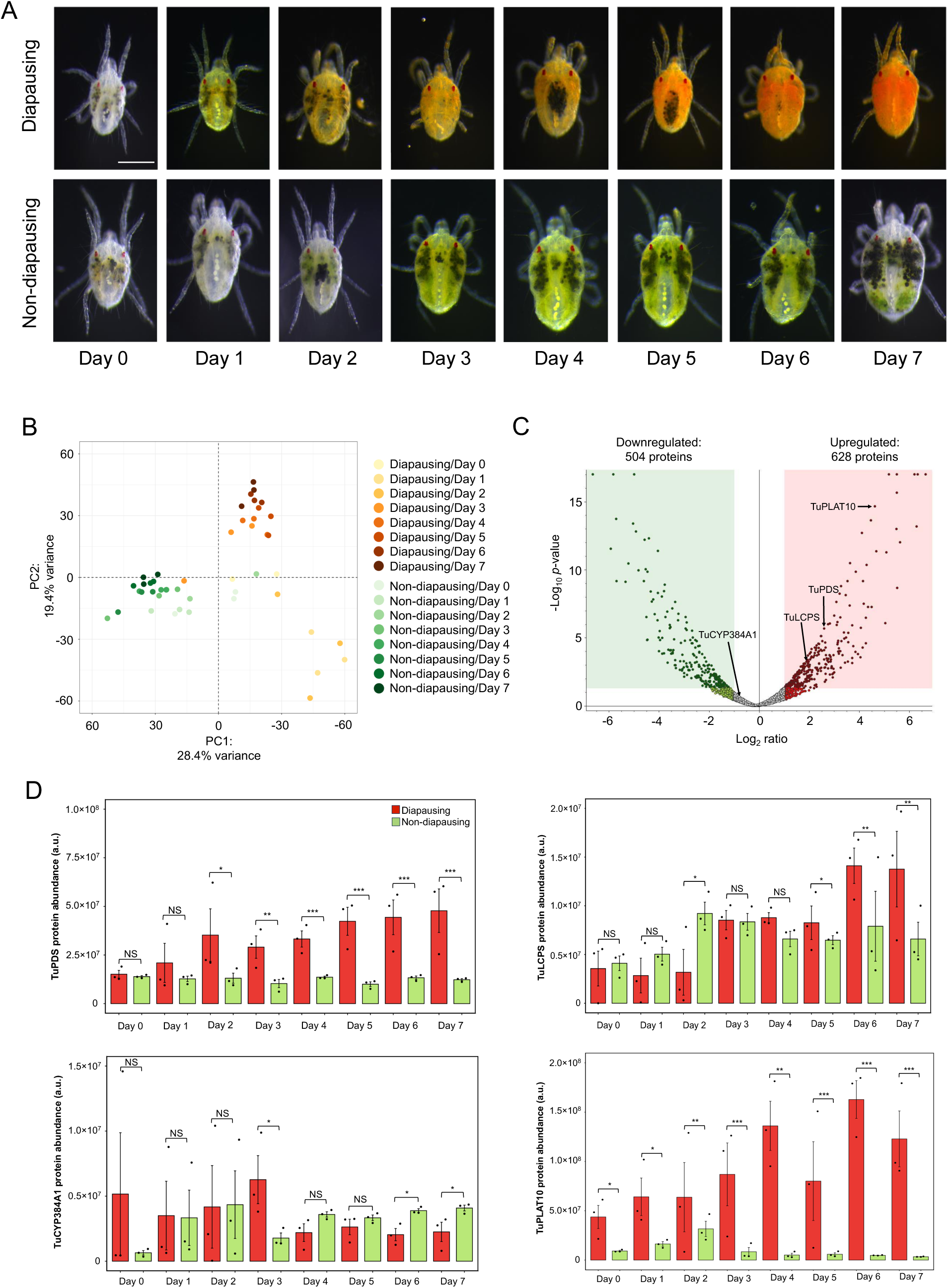
Proteomic analysis of diapausing and non-diapausing *Tetranychus urticae* females. (A) Representative images showing body color changes under diapause-inducing (LD 8:16) and non-diapause (reproduction)-inducing (LD 16:8) photoperiods at 18 °C across the 7-day period following the final molt to adulthood. Diapausing females progressively develop a uniform orange pigmentation, whereas non-diapausing females acquire a yellowish green pigmentation accompanying ovarian development. Scale bar, 0.1 mm. (B) Principal component analysis (PCA) of LC-MS/MS proteomic data. Each point represents a biological replicate. (C) Volcano plot of differentially expressed proteins between diapausing and non-diapausing females at day 7. The *x*-axis shows log_2_ fold change (FC) between diapausing and non-diapausing females, and the *y*-axis shows −log_10_ of the *p*-value. Dark green, significantly downregulated (log_2_FC < −1, *p* < 0.05); light green, downregulated but not statistically significant; dark red, significantly upregulated (log_2_FC > 1, *p* < 0.05); light red, upregulated but not statistically significant; gray, no significant change. Carotenoid biosynthesis enzymes (TuPDS, phytoene desaturase, *tetur01g11270*; TuLCPS, lycopene cyclase/phytoene synthase, *tetur01g11260*; TuCYP384A1, cytochrome P450 family member CYP384A1, candidate carotenoid ketolase, *tetur38g00650*) and the single PLAT (Polycystin-1, Lipoxygenase, Alpha-toxin) domain protein TuPLAT10 (*tetur11g05720*) are labeled. (D) Time-course relative protein abundance profiles of TuPDS, TuLCPS, TuCYP384A1, and TuPLAT10 from day 0 to day 7 under diapause and non-diapause conditions. Data are mean ± standard error of the mean (SEM) (*n* = 3 biological replicates). Statistical significance was assessed using Student’s *t*-test. NS (not significant), *p* ≥ 0.05; \**p* < 0.05; \*\**p* < 0.01; \*\*\**p* < 0.001.

To characterize the proteomic changes associated with this phenotypic transition, principal component analysis (PCA) was performed on proteomic data collected from diapausing and non-diapausing females across the time course (Fig. 1B). PC1 clearly separated diapausing from non-diapausing samples. Non-diapausing samples shifted slightly along PC1 over the time course but remained confined to the left region of the PCA plot. In contrast, diapausing samples migrated progressively along both PC1 and PC2 over the time course. From day 0 to day 3, diapausing samples partially overlapped with the non-diapausing cluster, and from day 4 onward they formed a tight cluster in the upper-right quadrant of the plot.

In line with this PCA trajectory, time-course volcano plots from day 0 to day 6 further illustrated the proteomic changes accompanying diapause progression (Fig. S1). Even at day 0, a considerable number of proteins were differentially abundant between diapausing and non-diapausing females in both up- and downregulated regions. As diapause progressed, the upregulated region (red) showed an increasing abundance of proteins at higher log_2_ fold change values from day 3 onward, whereas the downregulated region (green) maintained a relatively stable distribution throughout the time course. At day 7, a total of 2,316 downregulated and 1,896 upregulated proteins were identified in diapausing females relative to non-diapausing females (Fig. 1C).

Among the labeled proteins, TuPDS and TuLCPS are carotenoid biosynthesis enzymes encoded by genes horizontally transferred from fungi to *T. urticae* (Altincicek et al., 2012; Bryon et al., 2017) and TuCYP384A1 is a candidate carotenoid ketolase implicated by genetic mapping of a keto-carotenoid-deficient mutant in the closely related species *T. kanzawai* (Wybouw et al., 2019). These three proteins were therefore expected to show abundance changes associated with the carotenoid-based body pigmentation of diapausing females. TuPDS appeared in the upregulated region from day 2 onward (Fig. S1), reached high statistical significance at day 7 (Fig. 1C), and progressively increased in diapausing females over the time course, with significant differences from non-diapausing females emerging at day 2 and reaching the highest level of significance from day 4 to day 7 (Fig. 1D). TuLCPS became prominently upregulated from day 5 (Fig. S1) and remained markedly upregulated at day 7 (Fig. 1C). TuCYP384A1, in contrast, remained near the center of the volcano plot throughout day 0 to day 6 (Fig. S1) but became slightly downregulated by day 7 (Fig. 1C); consistently, its temporal abundance profile showed a transient upregulation at day 3, followed by a shift toward downregulation at days 6 and 7 (Fig. 1D). TuPLAT10, a 180-amino-acid single PLAT domain protein with no previously characterized role in diapause, was already upregulated at day 0 in the volcano plots and remained stably upregulated thereafter (Fig. S1). It was among the most strongly upregulated proteins at day 7 (Fig. 1C) and was significantly more abundant in diapausing than in non-diapausing females from day 0 onward, with this difference maintained throughout the 7-day time course (Fig. 1D). Of the 12 TuPLAT family members quantified in the proteomic analysis, only TuPLAT10 showed a statistically significant difference in abundance between diapausing and non-diapausing females at day 7 (Fig. S2).

### 3.2. RNAi disrupts diapause body pigmentation and carotenoid accumulation

To assess the functional roles of the four candidate proteins, RNAi-mediated silencing was performed in *T. urticae* females reared under diapause-inducing conditions using ds*TuPDS*, ds*TuLCPS*, ds*TuCYP384A1*, and ds*TuPLAT10*, with dsNC serving as the negative control. Body pigmentation phenotypes were classified into three categories: orange (typical diapause), yellow (intermediate), and green (reduced diapause-associated pigmentation; Fig. 2A). In the dsNC control, all individuals (100%, *n* = 128) developed the typical orange diapause pigmentation. ds*TuLCPS* treatment resulted predominantly in yellow individuals (78.8%, *n* = 119), with the remaining 21.2% orange individuals. In contrast, almost half of the individuals had green pigmentation in the ds*TuPDS* (44.4%, *n* = 115), ds*TuCYP384A1* (49.9%, *n* = 123), and ds*TuPLAT10* (51.3%, *n* = 122) treatments, together with yellow individuals (35.4%, 29.5%, and 17.7%, respectively) and reduced proportions of orange individuals (20.2%, 20.6%, and 31.0%, respectively). All four RNAi treatments differed significantly from the dsNC control (Fisher’s exact test, *p* < 0.001). Normal diapausing females retained the orange body color and well-defined red eye spots (Fig. S3A) and normal non-diapausing females showed the green body color with similarly intact red eye spots (Fig. S3B), whereas green-bodied ds*TuPLAT10*-treated individuals had reduced eye spot pigmentation, with several individuals showing partial or complete loss of the eye spots (Fig. S3C–F).

**Fig. 2.**
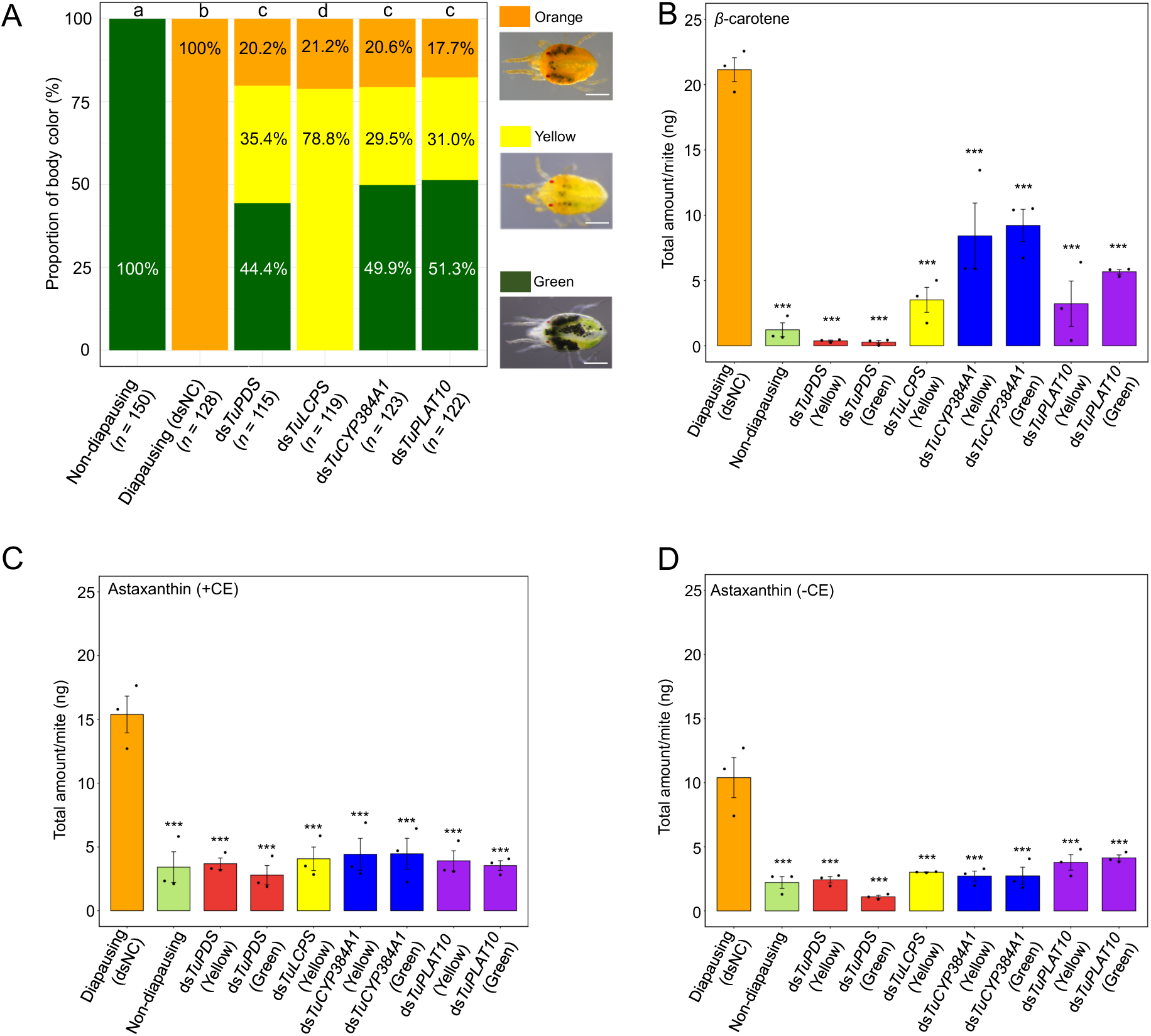
Body coloration phenotypes and carotenoid contents following RNAi-mediated gene silencing in *Tetranychus urticae* females reared under diapause-inducing conditions. (A) Representative images of the three body color phenotypes scored after dsRNA treatment: orange (typical diapause phenotype with high carotenoid accumulation), yellow (intermediate, altered carotenoid composition), and green (reduced diapause-associated carotenoid accumulation). Scale bars, 0.1 mm. The stacked bar chart represents the proportion of individuals classified as orange, yellow, or green within each treatment group following dsRNA treatments targeting the *TuPDS* (*tetur01g11270*), *TuLCPS* (*tetur01g11260*), *TuCYP384A1* (*tetur38g00650*), or *TuPLAT10* (*tetur11g05720*) genes (ds*TuPDS*, ds*TuLCPS*, ds*TuCYP384A1*, or ds*TuPLAT10*, respectively), compared with the dsRNA treatment targeting an intergenic region used as a negative control (dsNC). Different letters above the bars indicate significant differences among treatments (Fisher’s exact test, *p* < 0.001); *n*, total number of mites scored. Of females classified by body coloration phenotype, the body-color subgroups containing sufficient individuals were subjected to high-performance liquid chromatography (HPLC) analysis: yellow only for ds*TuLCPS*; yellow and green for ds*TuPDS*, ds*TuCYP384A1*, and ds*TuPLAT10*. dsNC females (all orange) and non-diapausing females (all green) were each analyzed as a single group. HPLC analysis for quantifying the contents of (B) β-carotene and (C, D) astaxanthin in dsRNA-treated mites and non-diapausing mites. (C) Cholesterol esterase treatment (+CE) hydrolyses esterified astaxanthin into its free forms. In (D), −CE indicates samples without esterase treatment (free astaxanthin only). Data are mean ± standard error of the mean (SEM), with individual dots indicating biological replicates. Statistical significance relative to dsNC was determined using Dunnett’s test (*p* < 0.001).

To examine the carotenoid composition underlying these phenotypes, the contents of β-carotene and astaxanthin were quantified by HPLC. For each RNAi treatment, individuals were stratified by their body coloration phenotype (Fig. 2A), and the subgroups containing sufficient individuals were subjected to HPLC analysis: yellow only for the ds*TuLCPS* treatment and both yellow and green for the ds*TuPDS*, ds*TuCYP384A1*, and ds*TuPLAT10* treatments. dsNC-treated females (all orange) and non-diapausing females (all green) were each analyzed as single groups. The β-carotene content was highest in dsNC-treated females and was substantially reduced in non-diapausing controls and in all RNAi color subgroups (Dunnett’s test, *p* < 0.001 vs. dsNC; Fig. 2B). Among these subgroups, ds*TuPDS*-treated females (both yellow and green) showed the most pronounced reductions in β-carotene content. Astaxanthin content was then quantified in two complementary ways: with cholesterol esterase treatment (+CE; Fig. 2C), which results in the hydrolysis of astaxanthin esters to the free form and therefore yields the sum of free and esterified astaxanthin, and without CE treatment (−CE; Fig. 2D), which detects only the originally non-esterified (free) astaxanthin pool. Total astaxanthin (+CE) and free astaxanthin (−CE) showed similar overall patterns, with diapausing dsNC-treated females having the highest values and all RNAi color subgroups and non-diapausing females showing significantly reduced levels (Dunnett’s test, *p* < 0.001 vs. dsNC). The difference between the +CE and −CE values, which corresponds to the amount of astaxanthin accumulated as fatty acid esters, was largest in dsNC-treated females and was diminished in all RNAi subgroups; this reduction was particularly pronounced in ds*TuPLAT10*-treated females (yellow and green subgroups).

### 3.3. *TuPLAT10* silencing restores reproduction and selectively reduces TuPDS and TuLCPS

To examine whether silencing of any of the four candidate genes affected the reproductive arrest characteristic of diapause, egg production was monitored over a 6-day period following dsRNA treatment (Fig. 3). Females treated with dsNC, ds*TuPDS*, ds*TuLCPS*, and ds*TuCYP384A1* produced no eggs during the observation period, consistent with maintained reproductive diapause. In contrast, ds*TuPLAT10*-treated females resumed oviposition (Dunn’s multiple comparison test, *p* < 0.001). Among the four candidates, only ds*TuPLAT10* was thus capable of breaking the reproductive arrest of diapause. Adult survival over a 12-day period was also monitored and showed significant differences among treatments (log-rank test with Bonferroni correction, *p* < 0.001; Fig. S4).

**Fig. 3.**
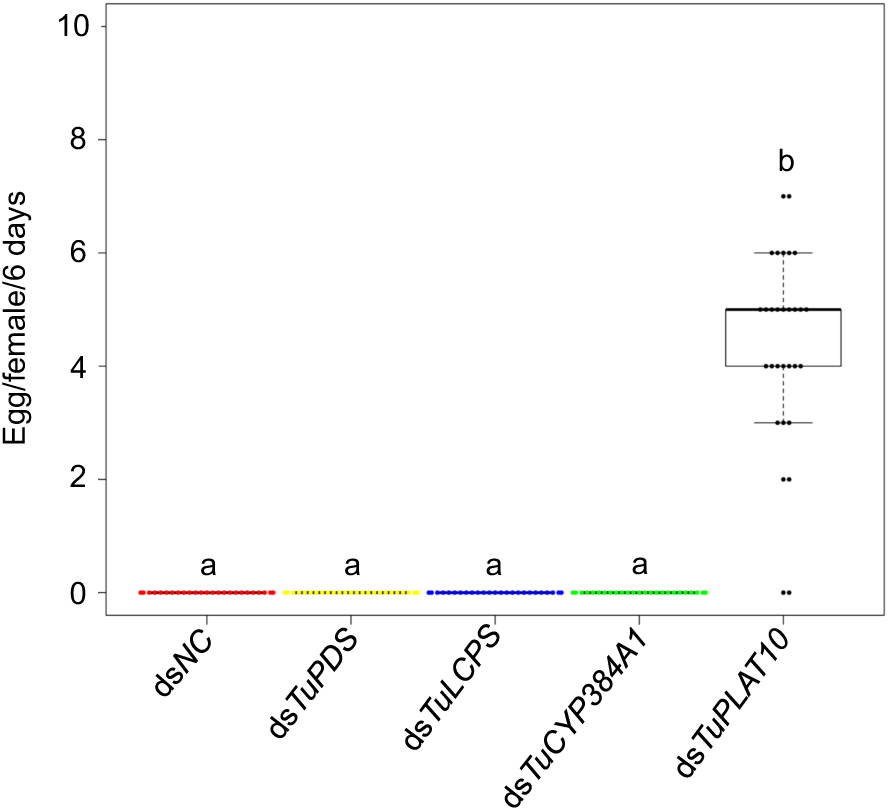
Fecundity of *Tetranychus urticae* females following RNAi-mediated gene silencing under diapause-inducing conditions. Egg production per surviving female per 6 days after adult emergence; dsNC, negative control. Only ds*TuPLAT10*-treated females oviposited, whereas no eggs were produced in the other treatment groups during the observation period. Different letters above the boxplots indicate significant differences (Dunn’s multiple comparison test, *p* < 0.001; *n* = 30).

The marked reduction in TuPLAT10 abundance in ds*TuPLAT10*-treated females compared with dsNC controls (Dunn’s multiple comparison test, *p* < 0.05), reaching levels comparable to those in non-diapausing controls, confirmed effective RNAi-mediated silencing of the target gene at the protein level (Fig. 4A). Despite the absence of sequence similarity between ds*TuPLAT10* and any other carotenoid biosynthesis gene, TuPDS abundance was significantly lower in ds*TuPLAT10*-treated females than in both dsNC and non-diapausing controls (Fig. 4B). TuLCPS abundance showed a similar pattern, differing significantly from that in both dsNC (which showed the highest abundance) and non-diapausing females (Fig. 4C). In contrast, TuCYP384A1 abundance did not differ significantly among dsNC, non-diapausing, and ds*TuPLAT10*-treated females (Fig. 4D). These results show that silencing of the *TuPLAT10* gene selectively reduced TuPDS and TuLCPS at the protein level without affecting TuCYP384A1.

**Fig. 4.**
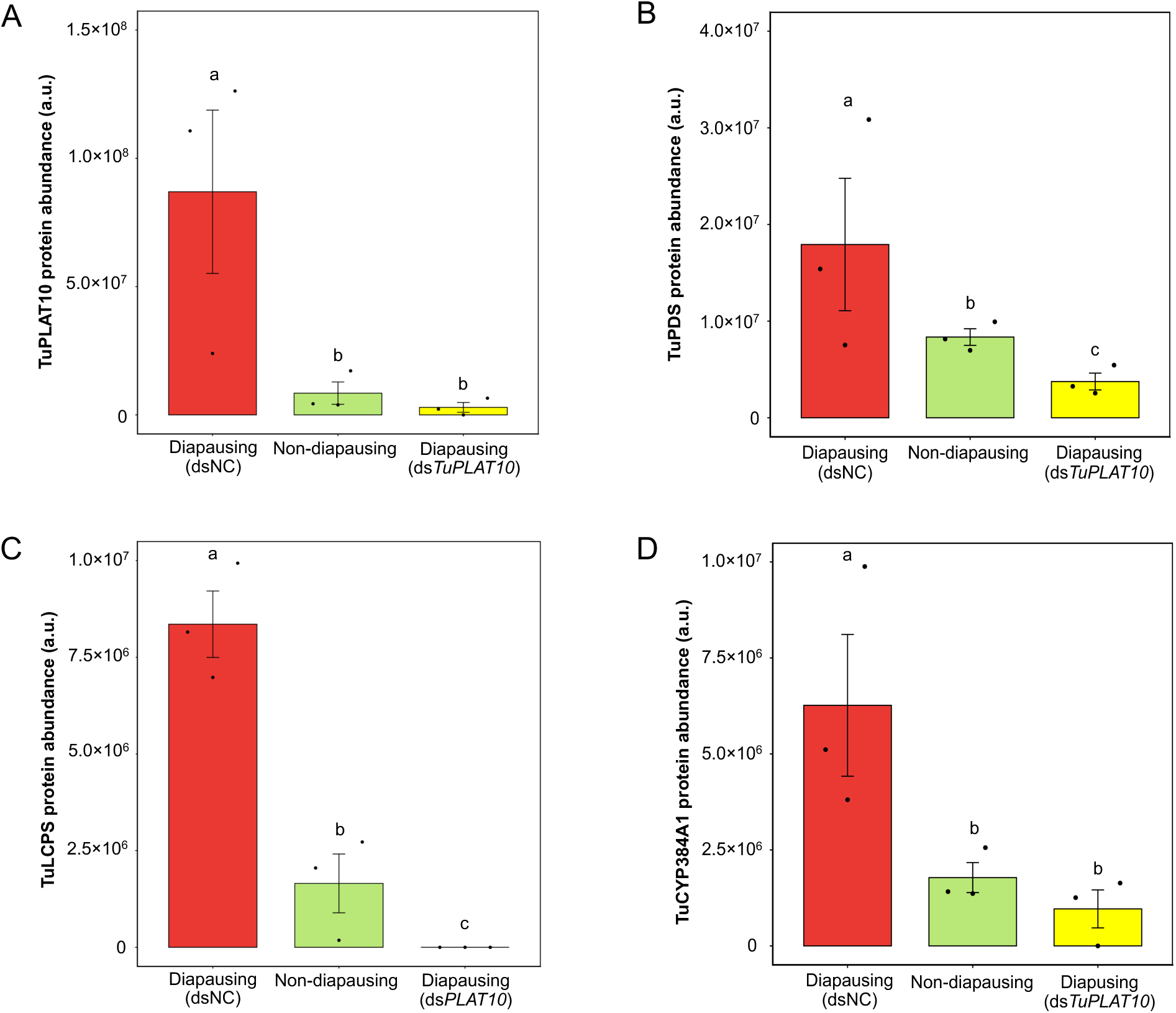
Effects of RNAi-mediated silencing of the *TuPLAT10* (*tetur11g05720*) gene on the protein abundances of (A) TuPLAT10, (B) TuPDS (phytoene desaturase, *tetur01g11270*), (C) TuLCPS (lycopene cyclase/phytoene synthase, *tetur01g11260*), and (D) TuCYP384A1 (cytochrome P450 family member CYP384A1, candidate carotenoid ketolase, *tetur38g00650*) in diapausing *Tetranychus urticae* females. Data are mean ± standard deviation (SD) for the negative control (dsNC), the control of non-diapausing females, and ds*TuPLAT10* treatment groups. Different letters above bars indicate significant differences (Dunn’s multiple comparison test, *p* < 0.05).

## 4. Discussion

The photoperiodic diapause of *T. urticae* females was first described by Bondarenko (1950) and has since been studied as a model of arthropod photoperiodism (Goto, 2016). The diapause phenotype combines reproductive arrest and orange body pigmentation, but how these two traits are molecularly coupled has remained an open question. The present study combined comparative proteomics between diapausing and non-diapausing females with RNAi-based functional analysis of candidate genes. RNAi suppression of the carotenoid biosynthesis enzymes TuPDS and TuLCPS markedly reduced both orange body coloration and the accumulation of β-carotene and astaxanthin (Fig. 2), providing functional support for HGT-derived carotenoid biosynthesis enzymes as essential factors of the diapause phenotype. The proteomic analysis additionally identified a single PLAT domain protein, TuPLAT10 (*tetur11g05720*), as a novel candidate regulator: ds*TuPLAT10* treatment simultaneously suppressed orange body coloration, restored oviposition under diapause-inducing conditions (Fig. 3), and selectively reduced TuPDS and TuLCPS protein levels without affecting the candidate ketolase TuCYP384A1 (Fig. 4). We propose that TuPLAT10 functions as a molecular switch that, in response to photoperiodic information, redirects the fate of fatty acids between egg production and pigment accumulation, and we develop this idea below as the phosphatidic acid (PA) allocation model (Fig. 5).

**Fig. 5.**
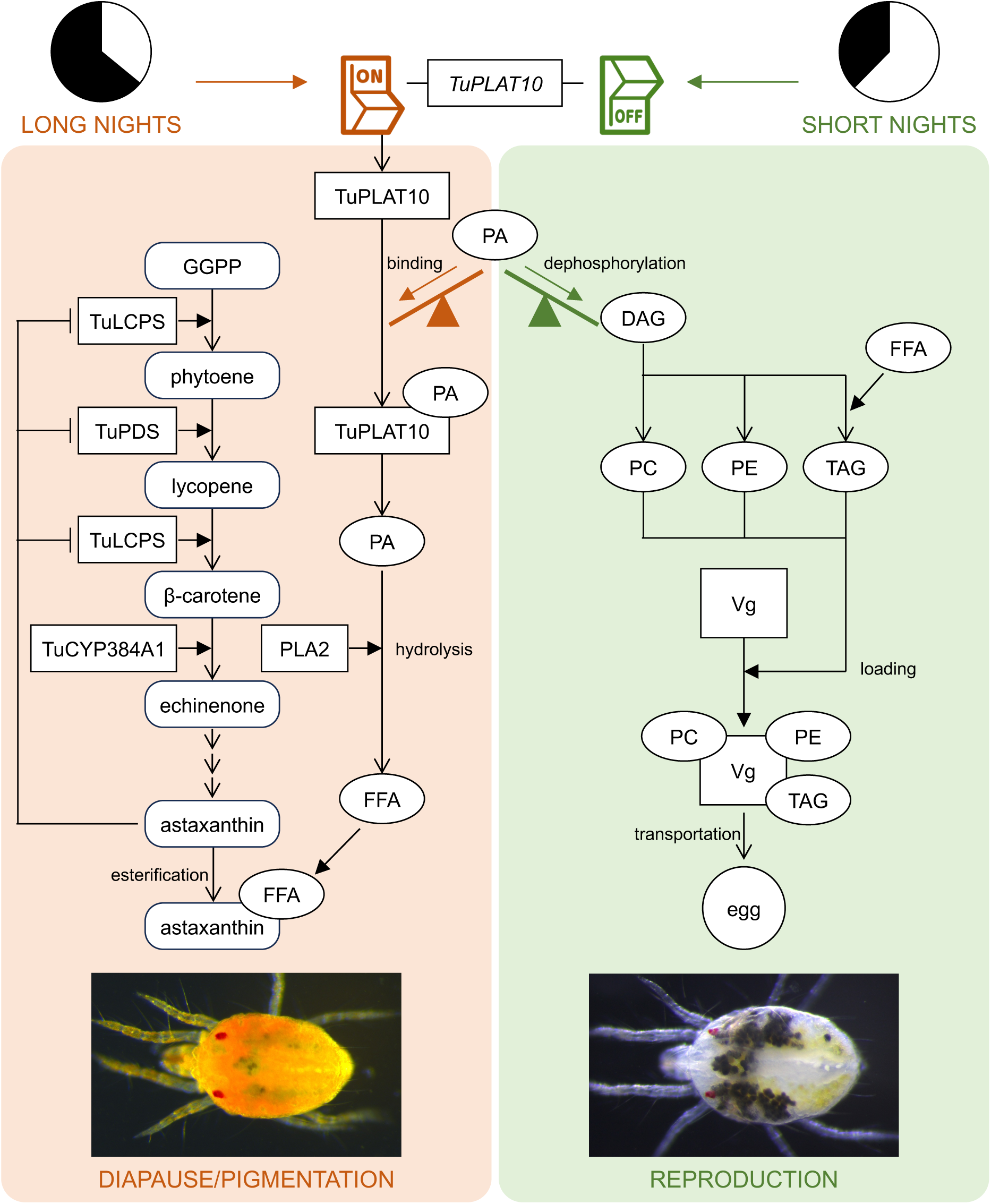
Proposed phosphatidic acid (PA) allocation model in which TuPLAT10 couples reproductive arrest and carotenoid pigmentation during diapause in *Tetranychus urticae* females. In midgut epithelial cells, PA sits at a central branch point of glycerolipid metabolism, and its allocation between two competing fates is depicted as a balance. Photoperiod regulates this balance through the expression of TuPLAT10 (*tetur11g05720*), a single PLAT domain protein predicted to function as a non-catalytic, membrane-bound PA-binding carrier. Under long-night photoperiods (left, diapause and pigmentation state), *TuPLAT10* is expressed (switched ON), and the encoded TuPLAT10 protein binds and sequesters PA (binding). The bound PA is then hydrolyzed by phospholipase A2 (PLA2) to release free fatty acids (FFAs) (hydrolysis), which are channeled into the esterification of astaxanthin (esterification). β-carotene is endogenously biosynthesized *de novo* from geranylgeranyl diphosphate (GGPP) through three enzymatic steps catalyzed by two enzymes encoded by horizontally transferred genes of fungal origin: the bifunctional lycopene cyclase/phytoene synthase TuLCPS (*tetur01g11260*; catalyzing both the GGPP to phytoene and the lycopene to β-carotene steps) and phytoene desaturase TuPDS (*tetur01g11270*; converting phytoene to lycopene). β-carotene is then converted via echinenone to astaxanthin by the candidate carotenoid ketolase TuCYP384A1 (*tetur38g00650*). Free astaxanthin is then esterified with the FFAs derived from PA hydrolysis to form astaxanthin fatty acid esters. The resulting astaxanthin fatty acid esters accumulate in lipid droplets of the midgut epithelial cells and produce the orange body pigmentation characteristic of diapausing females. In parallel, vitellogenin (Vg) synthesis is suppressed and lipid supply to the adjacent ovaries is shut off, resulting in reproductive arrest. T-bars extending from free astaxanthin to TuLCPS and TuPDS denote a tentative feedback inhibition of upstream carotenoid biosynthesis enzymes by free (non-esterified) astaxanthin. Under short-night photoperiods (right, reproductive state), *TuPLAT10* is switched OFF, and PA is no longer sequestered. Instead, PA is converted to diacylglycerol (DAG) by a PA phosphatase (dephosphorylation), and DAG enters the conventional Kennedy pathway to be incorporated into the membrane phospholipids phosphatidylcholine (PC) and phosphatidylethanolamine (PE) and into storage triacylglycerol (TAG); FFAs are also incorporated into TAG. These lipid resources are loaded onto Vg (loading) and supplied via Vg-mediated transport to the adjacent ovaries to support oogenesis (right photograph), enabling continuous egg production. Photoperiod-dependent expression of TuPLAT10 thus partitions PA between astaxanthin esterification (long nights) and Vg-mediated lipid supply for the yolk (short nights).

The PLAT domain is a β-sandwich module involved in lipid binding and protein–protein interactions (Bateman and Sandford, 1999). Single PLAT domain proteins (<200 amino acids, one PLAT domain only) were until recently considered plant-specific and have been collectively termed the PLAT-plant-stress protein family, contributing to abiotic stress tolerance and biotic defense (Hyun et al., 2014). Snoeck et al. (2018) reported that proteins of this family also occur in tetranychid mites; the genome of the polyphagous *T. urticae* contains 21 such genes, a marked family expansion compared with closely related oligophagous species. TuPLAT10 and its genomic neighbor TuPLAT11 (*tetur11g05730*) display nearly identical expression patterns under both host-plant adaptation (Snoeck et al., 2018) and photoperiodic responses (this study), suggesting a common regulatory mechanism responsive to environmental signals. We chose TuPLAT10, which exhibited the largest differential abundance in our proteome (Fig. S2), as the target of functional analysis.

Among plant single PLAT domain proteins, AtPLAT1, also known as PLAFP (Phloem Lipid-Associated Family Protein), in *Arabidopsis thaliana* (L.) Heynh. binds PA specifically and functions as a soluble carrier (Barbaglia et al., 2016; Kulke et al., 2024). Although sequence similarity to plant PLAFP is lacking, TuPLAT10 (180 amino acids) shares the conserved β-sandwich architecture, suggesting that PA binding may be conserved between the two proteins. InterPro analysis indicates that TuPLAT10 contains a PLAT/LH2 domain (IPR001024) as its sole functional domain and lacks any catalytic domains characteristic of lipoxygenases or lipases; an N-terminal signal peptide and a single transmembrane helix are also predicted. TuPLAT10 is thus most likely not a catalytic enzyme but a membrane-anchored PA-binding carrier, performing a PLAFP-like function in a membrane-tethered form.

A PA carrier could coordinately regulate diapause and reproduction in *T. urticae* because of the compact body plan of this species, in which multiple metabolic functions are consolidated within the midgut epithelium. Unlike other arthropods, *T. urticae* lacks both a developed hemolymph circulation and an independent fat body (Blauvelt, 1945; Mothes-Wagner, 1982; Bensoussan et al., 2018). Midgut epithelial cells therefore integrate digestion and absorption with lipid storage, detoxification, and vitellogenin synthesis, functions that in insects are largely fulfilled by the fat body. Because the ventral midgut epithelium lies directly adjacent to the ovaries and developing eggs (Bensoussan et al., 2018), and because astaxanthin fatty acid esters that produce the orange body color also accumulate in lipid droplets of the same midgut epithelial cells (McEnroe, 1970; Kawaguchi et al., 2016), lipid metabolism, carotenoid metabolism, and reproduction-related protein synthesis are all integrated within the midgut epithelium.

In this context, PA stands at a central branch point of glycerolipid metabolism: It can flow via diacylglycerol into membrane phospholipids (phosphatidylcholine and phosphatidylethanolamine) and storage triacylglycerol through the Kennedy pathway (Kennedy, 1961), or its constituent fatty acids can be released by lipases and phospholipases into a free fatty acid pool that is used for the esterification of astaxanthin (Kawaguchi et al., 2016). Our PA allocation model (Fig. 5) proposes that TuPLAT10 governs this branch point: in the diapause state (high TuPLAT10), PA-derived free fatty acids are selectively channeled toward astaxanthin esterification, producing the orange body color in midgut lipid droplets, while vitellogenin synthesis is suppressed (Kawakami et al., 2009) and lipid supply to the ovaries is shut off, leading to reproductive arrest; in the reproductive state (low TuPLAT10), PA flows through the Kennedy pathway into vitellogenin-bound lipids that are supplied to the adjacent ovaries for egg formation. The difference between +CE and −CE astaxanthin values, which corresponds to the esterified astaxanthin pool, was somewhat smaller in ds*TuPLAT10*-treated females than in dsNC controls (Fig. 2C, D). This pattern suggests a reduced free fatty acid supply for esterification under TuPLAT10 suppression.

In addition to reducing the level of TuPLAT10 itself (Fig. 4A), ds*TuPLAT10* treatment significantly reduced TuPDS and TuLCPS abundances (Fig. 4B, C). Because there is no sequence similarity between ds*TuPLAT10* and these carotenoid biosynthesis genes, this reduction is unlikely to reflect an off-target RNAi effect. One possible explanation is feedback inhibition of upstream carotenoid biosynthesis by free astaxanthin. In the astaxanthin-producing green microalga *Haematococcus pluvialis* Flotow, astaxanthin esterification has been proposed to drive carotenoid flux by relieving such feedback inhibition (Chen et al., 2015; Basiony et al., 2022). Impaired esterification upon TuPLAT10 suppression could lead to free astaxanthin accumulation and consequent feedback reduction of upstream enzymes such as TuPDS and TuLCPS, thereby limiting carotenoid production and accounting for the suppressed orange body pigmentation in ds*TuPLAT10*-treated females (Fig. 2A). However, direct evidence for this potential mechanism in *T. urticae* is lacking and remains a topic for future biochemical study.

The photoperiodic system in arthropods consists of four functional units: a photoreceptive input, a photoperiodic clock, a photoperiodic counter, and a neuroendocrine output that triggers the phenotypic shift (Goto, 2016). In insects, cessation of juvenile hormone (JH) secretion at the neuroendocrine output corresponds to adult diapause, and topical application of a JH analog terminates diapause and induces ovarian development (Numata and Hidaka, 1984). In crustaceans, methyl farnesoate (MF) plays an analogous role to JH (Laufer et al., 1993).

Chelicerates lack JH and rely on MF as the JH-equivalent hormone (Qu et al., 2018). The *T. urticae* genome lacks the gene encoding the JH-epoxidating CYP15A1 but contains the gene encoding a farnesoic acid *O*-methyltransferase (FAMeT)-like protein (*tetur30g00780*), an enzyme of MF biosynthesis (Grbić et al., 2011). A direct role of MF in *T. urticae* diapause has yet to be experimentally demonstrated, but its possible involvement has been postulated (Goto, 2016). The resumption of oviposition under diapause-inducing conditions caused by ds*TuPLAT10* is functionally analogous to JH analog–induced diapause termination in insects (Numata and Hidaka, 1984) and may reflect a corresponding role of MF in *T. urticae*. Within this framework, TuPLAT10 may operate downstream of the proposed MF-based neuroendocrine output as an effector that translates such a signal into the metabolic event of PA partitioning. However, we cannot rule out a parallel and independent role for TuPLAT10, and the hierarchical relationship between TuPLAT10 and the endocrine system remains a question for future work.

Based on our findings, we have proposed a working hypothesis that integrates reproductive arrest and carotenoid pigmentation along a common metabolic axis of PA partitioning mediated by TuPLAT10 (Fig. 5). Priorities for future research include (i) lipidomic analysis focused on PA in ds*TuPLAT10*-treated females; (ii) characterization of PA binding by recombinant TuPLAT10 and elucidation of its tissue localization; (iii) biochemical verification of whether free astaxanthin directly inhibits TuPDS or TuLCPS; and (iv) clarification of the hierarchical relationship between TuPLAT10 and the endocrine system through MF administration or RNAi of the *FAMeT* gene. These studies will clarify how the unique anatomy of spider mites shapes the integration of lipid, carotenoid, and reproductive metabolism and may inform the development of diapause-targeted pest control.

## CRediT authorship contribution statement

**Rismayani:** Writing – original draft, Writing – review & editing, Visualization, Validation, Software, Methodology, Investigation, Formal analysis, Conceptualization. **Kanae Sai:** Data curation, Investigation. **Tomohiro Ohsako:** Data curation, Investigation. **Murata Kohyoh:** Data curation, Investigation. **Yuka Arai:** Data curation, Investigation. **Naoki Takeda:** Data curation, Investigation. **Masanobu Yamamoto:** Data curation, Investigation. **Rika Umemiya-Shirafuji**: Funding acquisition, Writing – review & editing. **Takeshi Suzuki:** Supervision, Funding acquisition, Project administration, Resources, Methodology, Writing – review & editing, Formal analysis, Conceptualization.

## Disclosure

The authors declare no conflicts of interest.

## Acknowledgments

The authors would like to thank Dr. T. Gotoh (Ibaraki University, Ibaraki, Japan) and Drs. M. Grbić and V. Grbić (The University of Western Ontario, London, ON, Canada) for providing the mite populations used in this study. This work was supported by the SDGs Global Leader Program, Japan International Cooperation Agency (JICA). This work was also supported in part by JSPS KAKENHI grants 16K18661, 18H02203, 21H02193, and 24K21256, as well as the Cabinet Office, Government of Japan, Cross-ministerial Moonshot Agriculture, Forestry and Fisheries Research and Development Program, “Technologies for Smart Bio-industry and Agriculture” (JPJ009237; funding agency: Bio-oriented Technology Research Advancement Institution), awarded to TS. This work was also supported in part by Cooperative Research Grants (2024joint-9, 2025joint-1, and 2026joint-5) of the National Research Center for Protozoan Diseases, Obihiro University of Agriculture and Veterinary Medicine, awarded to RUS and TS.

## Notes

### Competing Interest Statement

The authors have declared no competing interest.

### Summary of Updates

Reference list updated with corrected citation details.

