## Supplemental Figure 1 for "A single PLAT domain protein couples reproductive arrest and carotenoid pigmentation during diapause in the two-spotted spider mite, *Tetranychus urticae* Koch"

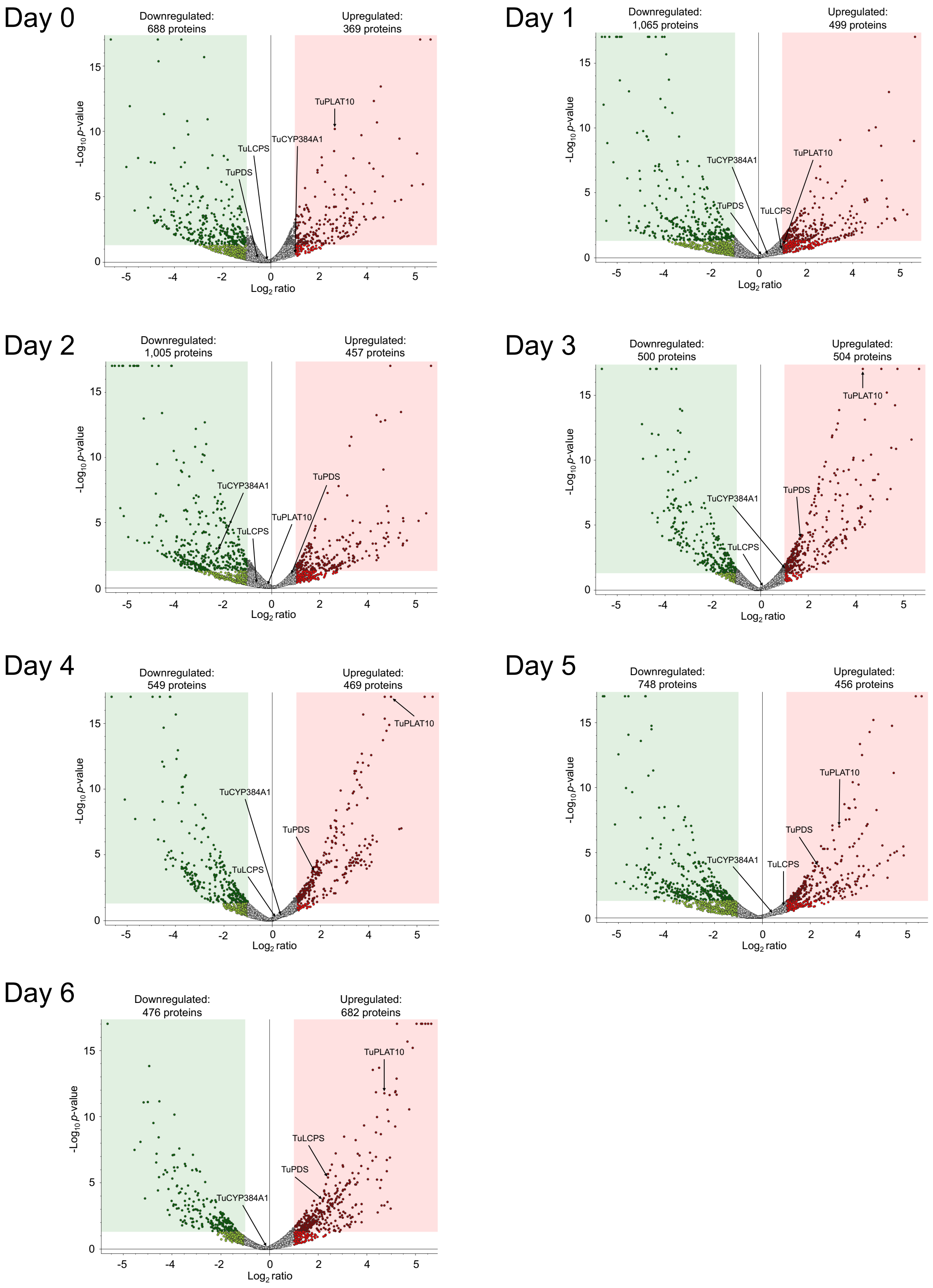


**Fig. S1.** Time-course volcano plots of proteomic differences between diapausing and non-diapausing *Tetranychus urticae* females from day 0 to day 6 after the final molt. The *x*-axes show

$\log_2$ fold change (diapausing/non-diapausing) and the *y*-axes show $-\log_{10}$ (*p*-value). The green and red shaded regions indicate significantly downregulated and upregulated proteins, respectively. The latter increases progressively with diapause progression. Carotenoid biosynthesis enzymes (TuPDS, phytoene desaturase; TuLCPS, lycopene cyclase/phytoene synthase; TuCYP384A1, cytochrome P450 family member CYP384A1, candidate carotenoid ketolase), and the single PLAT (Polycystin-1, Lipoxygenase, Alpha-toxin) domain protein TuPLAT10 are labeled to indicate their dynamics across the time course.
