## Supplemental Figure 2 for "A single PLAT domain protein couples reproductive arrest and carotenoid pigmentation during diapause in the two-spotted spider mite, *Tetranychus urticae* Koch"

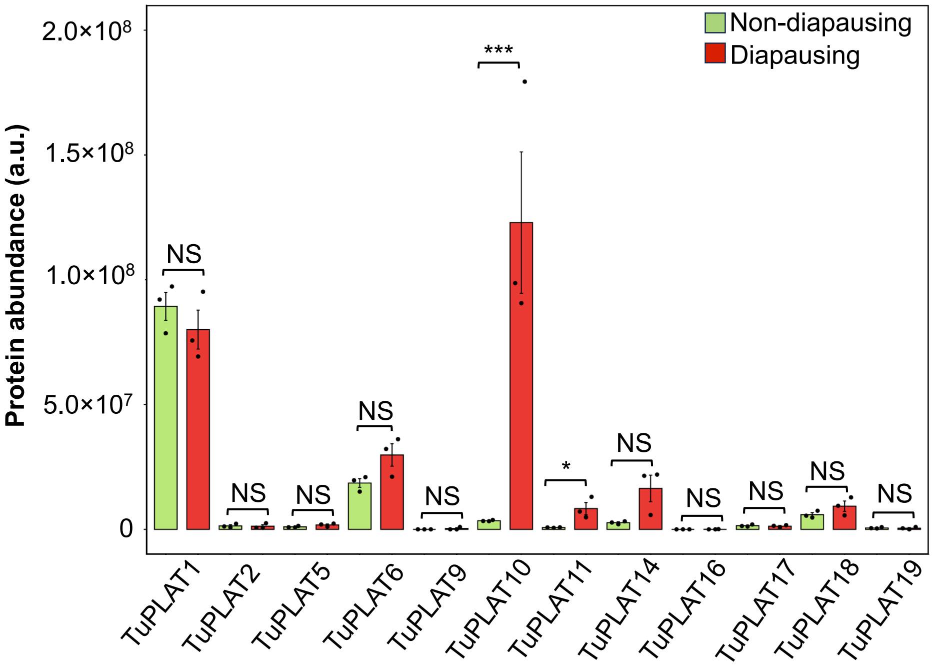


**Fig. S2.** Relative protein abundance of TuPLAT (PLAT [Polycystin-1, Lipoxygenase, Alpha-toxin] domain protein) family members in diapausing and non-diapausing *Tetranychus urticae* females at day 7 after the final molt. Bar graphs show the relative protein abundance of the 12 TuPLAT family members quantified in the proteomic analysis. Black dots represent individual biological replicates. Data are mean ± standard error of the mean (SEM). Among all TuPLAT family members detected, only TuPLAT10 showed a statistically significant difference between diapausing and non-diapausing females. NS (not significant), $p \geq 0.05$; *$p < 0.05$; ***$p < 0.001$ (Student's *t*-test).
