## Supplemental Figure 3 for "A single PLAT domain protein couples reproductive arrest and carotenoid pigmentation during diapause in the two-spotted spider mite, *Tetranychus urticae* Koch"

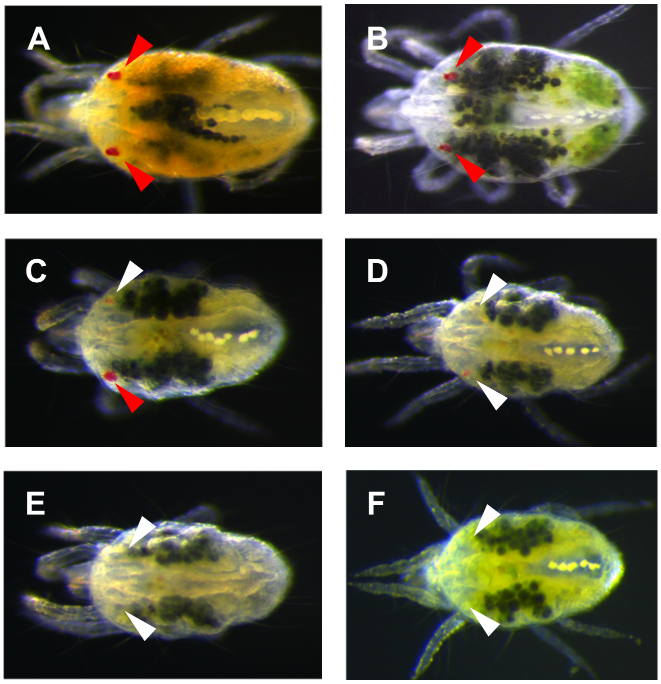


**Fig. S3.** Body and eye spot phenotypes following RNAi-mediated silencing of the *TuPLAT10* (single PLAT [Polycystin-1, Lipoxygenase, Alpha-toxin] domain protein 10) gene in adult *Tetranychus urticae* females. Representative images showing changes in body coloration and eye spot pigmentation. (A) Normal diapausing female with orange body color and well-defined red eye spot pigmentation (red arrowheads). (B) Normal non-diapausing female with green body color and well-defined red eye spot pigmentation (red arrowheads). (C) ds*TuPLAT10*-treated individual with green body coloration and well-defined red eye spot pigmentation on the one side (red arrowhead) but partial loss of eye spot pigmentation on the other side (white arrowhead). (D) ds*TuPLAT10*-treated individuals showing green body coloration with partial loss of eye spot pigmentation (white arrowheads). (E, F) ds*TuPLAT10*-treated individuals with green body coloration and a complete loss of eye spots. The reduction or loss of eye spot pigmentation in ds*TuPLAT10*-treated individuals is consistent with the disruption of carotenoid metabolism associated with *TuPLAT10* silencing, given that the red eye spots of *T. urticae* are themselves carotenoid-based.
