## Supplemental Figure 4 for "A single PLAT domain protein couples reproductive arrest and carotenoid pigmentation during diapause in the two-spotted spider mite, *Tetranychus urticae* Koch"

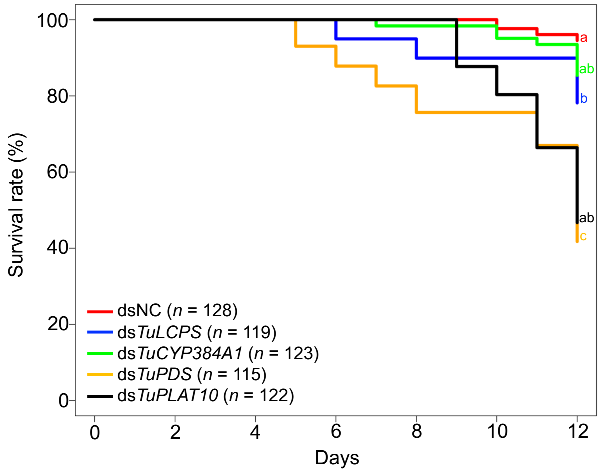


**Fig. S4**. Kaplan–Meier survival curves of *Tetranychus urticae* females following RNAi-mediated gene silencing under diapause-inducing conditions. Survival was observed for 12 days after adult emergence. dsNC, negative control. Significance of differences among treatments was determined by pairwise log-rank test with Bonferroni correction ($p < 0.001$). Different letters indicate statistically significant differences among groups.
