## Supplemental Table 1 for "A single PLAT domain protein couples reproductive arrest and carotenoid pigmentation during diapause in the two-spotted spider mite, *Tetranychus urticae* Koch"

**Table S1.** Primers for dsRNA synthesis. The T7 promoter sequence is underlined. NC: an intergenic region (Suzuki et al., 2017b) used as a negative control.

| Target | 5′-3′ Forward primer | 5′-3′ Reverse primer | Fragment size (bp) |
|---|---|---|---|
| *TuPDS* | TAATACGACTCACTATAGGGGTCCTGGCCCGTACAGTTTA | TAATACGACTCACTATAGGGCCCATTTGCTTTTCAATCGT | 625 |
| *TuLCPS* | TAATACGACTCACTATAGGGATTTGCTTCAAGGCCAAAAA | TAATACGACTCACTATAGGGCCAGAACAGGAGCTTGAACC | 635 |
| *TuCYP384A1* | TAATACGACTCACTATAGGGGCCTTTCAATGTCCATCCAT | TAATACGACTCACTATAGGGAAGCAAAAGGCAGGTAAGCA | 649 |
| *TuPLAT10* | TAATACGACTCACTATAGGGGTTCTACCATGGGCCCACAA | TAATACGACTCACTATAGGGCCACGTGCTGTTAATCTTACCA | 325 |
| NC | TAATACGACTCACTATAGGGCGACCCCATCAGGCTATTGA | TAATACGACTCACTATAGGGGCCCTCTCCTGGTTGTAAACTT | 382 |
